# Biologically grounded brain-on-chip model identifies selective topographic reorganization within hyperexcitable neural networks

**DOI:** 10.1101/2024.09.30.615880

**Authors:** Maxime Poinsot, Marine Dos Santos, Baptiste Marthy, Ana Borges-Correia, Eduardo Gascon-Gonzalo, Benoit Charlot, Maxime Cazorla

## Abstract

Connectomics has revolutionized our understanding of brain function by emphasizing the importance of neural networks and their topographical organization. Corticostriatal circuits, which play a critical role in cognition and emotion, follow a precise topographic architecture essential for integrating and processing cortical information within the basal ganglia. Disruptions to this connectivity are often implicated in neurodevelopmental and psychiatric disorders such as obsessive compulsive disorders, schizophrenia, epilepsy, and autism spectrum disorders. However, studying network disruptions *in vivo* presents significant challenges due to their intricate architecture and early developmental onset. To address this, we employed a brain-on-chip microfluidic platform to recreate a biologically relevant model of topographically organized corticostriatal networks. By mimicking the directional control of neuronal projections using Tesla valve-inspired microchannels, we demonstrate that genetic perturbations affecting neuronal excitability during development lead to selective alterations of local versus long-range network topology, resulting in the formation of new convergent nodes. This model offers critical insights into how early perturbations contribute to circuit-specific pathologies, providing a valuable tool for understanding neurodevelopmental disorders and advancing therapeutic strategies.

## INTRODUCTION

Connectomics, the study of brain connections, has driven breakthrough mapping initiatives such as the Human Connectome Project, the BRAIN Initiative, and the Blue Brain Project (Nowinski, 2017). Given its crucial role in high-order functions, cognition and emotion, the cerebral cortex has been a main focus of connectome research. Each cortical area sends descending projections that innervate subcortical structures, including the striatum, the primary input of to the basal ganglia. Corticostriatal projections are hierarchically organized and serially ordered, following a complex and precise topographic architecture, known to be instrumental in the transduction, integration, and processing of cortical information within the basal ganglia (Alexander et al., 1986; Parent and Hazrati, 1995; Graybiel, 2008). Anatomical studies across various species, including primates and rodents, have consistently demonstrated the presence of topographically organized corticostriatal pathways, highlighting integrative zones of convergence at the border of specific projection territory (Flaherty and Graybiel, 1991; Berendse et al., 1992; Hoffer and Alloway, 2001; Mailly et al., 2013; Averbeck et al., 2014; Hintiryan et al., 2016). According to the prevailing model, cortical output projections are functionally segregated into parallel cortico-basal ganglia-thalamo-cortical loops, each dedicated to the transmission and integration of specific types of information (sensorimotor, associative, or limbic) (Alexander et al., 1986; Haber, 2003; Oh et al., 2014).

The organizational principle of corticostriatal pathways has been instrumental in elucidating the pathophysiology of various neural disorders. Alterations in brain connectivity are often associated with disastrous cognitive and behavioral consequences in a wide array of neurodevelopmental and psychiatric diseases (Bullmore and Sporns, 2009; Crossley et al., 2014), including obsessive compulsive disorder (OCD) (Milad and Rauch, 2012; Ahmari et al., 2013), schizophrenia (Bullmore et al., 1997; Liu et al., 2008), epilepsy (Royer et al., 2022), and autism spectrum disorder (ASD) (Zeng et al., 2017). These dramatic neurological disorders, characterized by disrupted connectivity and activity in specific brain regions, often find their pathological origin during neurodevelopment (Shepherd, 2013; Nelson and Kreitzer, 2014). The formation and refinement of neural circuits typically occurs in-utero and during post-natal life, making early brain development a critical and vulnerable period for the establishment of brain networks. Genetic or external factors altering electrophysiological properties of neurons can affect axonal growth and synaptic connectivity, creating an excitation/inhibition imbalance in corticostriatal networks, linked with increased probability for neurodevelopmental disorders (Wolfram and Baines, 2013). For instance, genetic mutations in the inwardly rectifying potassium Kir2.1 channel gene altering neuronal excitability has been linked with dysfunctional cortical networks and is associated with a genetic risk for bipolar disorders, schizophrenia, mental retardation, epilepsy, and ASD (Sicca et al., 2011; Imbrici et al., 2013; Ambrosini et al., 2014; Binda et al., 2018). Therefore, understanding the neurodevelopmental origins of brain circuit topology in the healthy and diseased brain is essential, yet remains a significant challenge.

Recent progress in microfluidics and organ-on-chip technology has made it possible to develop unique platforms for creating *in vitro* models that approximate the complex environment of the brain (van der Meer and van den Berg, 2012; Neto et al., 2016). Brain-on-chip are state-of-the-art microfluidic devices used for a minimalistic reconstruction of neural circuits, in health and disease. Freedom offered by the design allows the creation of complex neuronal networks, using co-cultures and proper directional control of neuronal connections (Peyrin et al., 2011; Hasan and Berdichevsky, 2016; Honegger et al., 2016; Renault et al., 2016; Dauth et al., 2017; Gladkov et al., 2017; Holloway et al., 2019). Compartmentalization of neuronal cells and their neurites further allows independent control of each actor of the network (Taylor et al., 2005; Taylor et al., 2010; Moutaux et al., 2018a; Moutaux et al., 2018b; Virlogeux et al., 2018). Moreover, these microsystems offer unprecedented spatiotemporal resolution, enabling investigations of cellular and molecular mechanisms with high precision for the study of brain disorders and drug screening or pharmacological assays (Taylor et al., 2005; Yi et al., 2015; Nikolakopoulou et al., 2020; Holloway et al., 2021; Virlogeux et al., 2021; Miny et al., 2022).

Here we used a brain-on-chip microfluidic strategy to reconstitute a biologically grounded model of topographically organized corticostriatal circuits. We recapitulated the specific orientation of corticostriatal projections using Tesla valve-inspired asymmetric microchannels. By integrating morphological and functional analytic tools, we show that genetic perturbation of neuronal excitability during development profoundly destabilizes the topology of corticostriatal networks, creating new convergent nodes. This platform highlights how early developmental perturbations selectively affect the local and long-range architectural organization of the brain wiring diagram, offering new insights into circuitry-specific connectopathies.

## RESULTS

### Topographic reconstruction of corticostriatal pathways using microfluidic-based devices

We reconstructed the topographic organization of two segregated corticostriatal networks *in vitro* using a silicon polymer-based microfluidic device (**Figure 1a)**. We designed the microdevice using the open-access comprehensive corticostriatal projectome map developed by Hintiryan and colleagues (Hintiryan et al., 2016). This design emulates the architectural configuration of somatic sensorimotor and medial prefrontal cortical areas, which project to the ventrolateral and dorsomedial regions of the striatum, respectively (**Figure 1a)**. We selected these corticostriatal subnetworks due to the clear segregation of their terminal fields within the striatum and their specialized roles in processing and transferring sensory and executive information, respectively (Alexander et al., 1986). Additionally, the organization of these circuits is significantly altered in mouse models of disconnection syndromes, such as the zQ175 model of Huntington’s disease and the monoamine oxidase (MAO) A/B knockout model of autism-like behaviors (Hintiryan et al., 2016), making these two opposing pathways an ideal model for studying topographical rearrangement *in vitro*. The chip was configured to establish two parallel, opposing cortical chambers that project unidirectionally toward their respective striatal target areas, arbitrarily referred to here as dorsomedial (DM) and ventrolateral (VL) corticostriatal networks (**Figure 1b**). This arrangement also creates a central non-overlapping (NO) zone, where striatal neurons remain isolated from cortical axonal projections (**Figure 1b,c**). Each cortical chamber was seeded with a pool of rat primary cortical neurons, while the central postsynaptic target chamber was uniformly populated with striatal neurons (**Figure 1c**). The neuronal compartments are fluidically isolated and interconnected exclusively through a series of 500 μm-long microchannels. Given that cortical dendrites have a maximum growth length of approximately 450 μm (Taylor et al., 2005; Virlogeux et al., 2018), only axons from cortical neurons were allowed to extend through the microchannels into the striatal compartment. We further adapted the dimensions of the microchannels to optimize fluidic isolation and to reduce the number of axons in each channel, by reducing their width and height to 3 μm and 4 μm respectively (**Figure 1d**).

**Figure 1.**
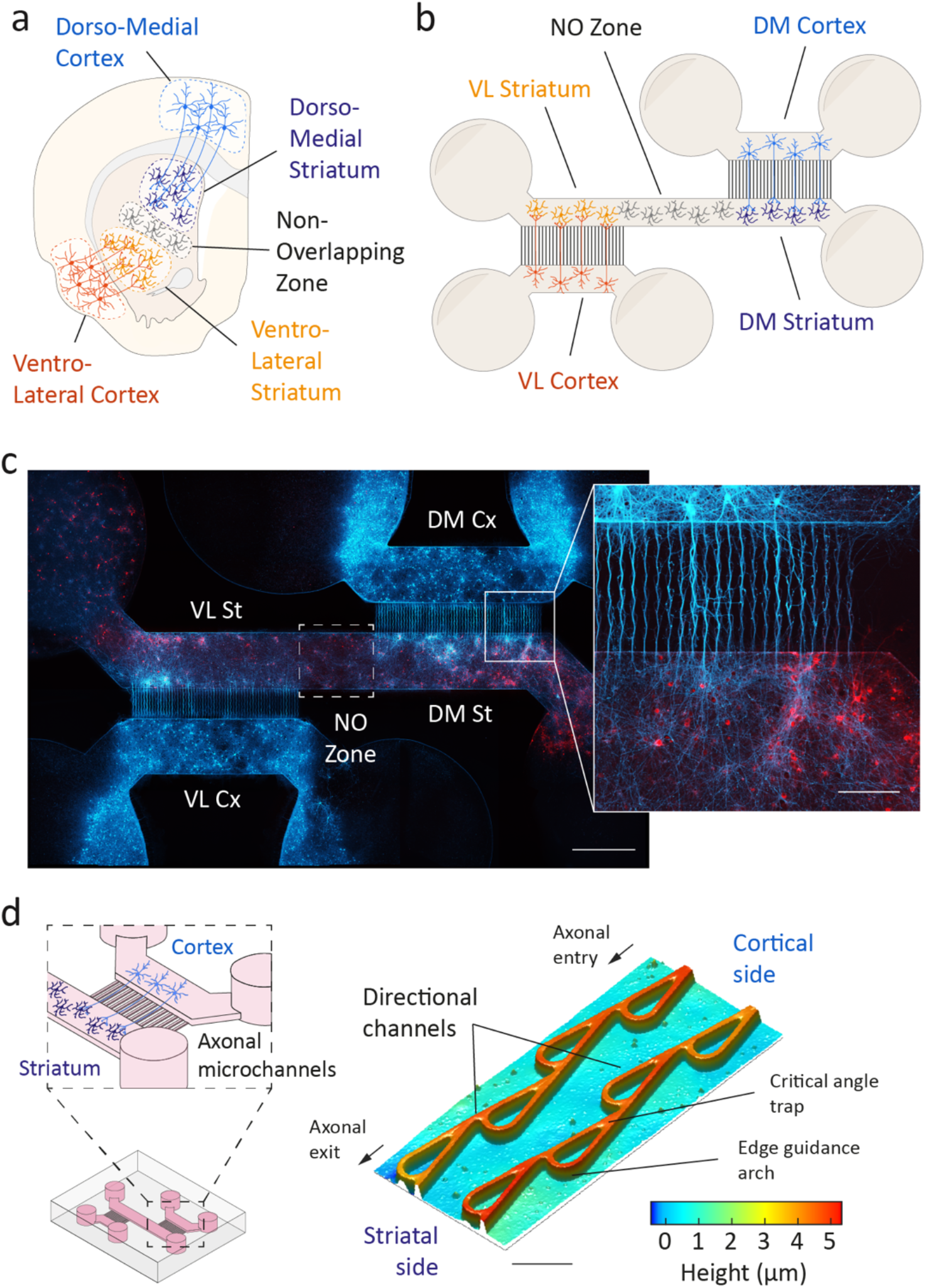
Biologically grounded model for the reconstruction of topographically segregated corticostriatal networks. **(a)** Schematic representation of dorsomedial and ventrolateral cortex regions (DM Cx, VL Cx) topographically projecting onto their respective striatal subregions (DM St, VL St) and separated by a non-overlapping (NO) zone. **(b)** Schematic of the 3-compartment microfluidic device allowing fluidic isolation of cortical and striatal chambers that are connected via unidirectional microchannels. An intermediate space devoid of cortical projections is inserted between both sets of microchannels in the striatal chamber to replicate the non-overlapping zone. Cylinders represent accessible wells for seeding neurons and perfusing liquids. **(c)** Mosaic reconstruction showing GFP-expressing cortical neurons (cyan) projecting through unidirectional microchannels onto mCherry-expressing striatal neurons. A non-overlapping (NO) zone devoid of cortical inputs is created between the two direct-projection fields (shown in inset). Scale bar, 1000 μm (Inset, 200 μm). **(d)** Tesla valve geometry was used to create unidirectional microchannels between cortical and striatal compartments. Phase contrast image of Tesla valve microchannel segment showing the critical angle and edge guidance constraints. Microchannel dimensions and geometries are color-coded. Scale bar, 30 μm.

### Tesla valve-inspired microchannels provide fluidic and axonal directionality

The primary challenge in conceiving physiologically-relevant models of segregated pathways is to control the unidirectionality of cortical axons projecting toward the striatal population, while preventing aberrant retrograde striato-cortical connections. To achieve this, we designed microchannels following the original geometry of valvular conduits proposed by Nikola Tesla in 1920 (Tesla, 1920) (**Figure 1d**). The unique architecture of the Tesla valve confers a key dual functionality advantage, including a robust diode effect for fluidic movement in channels and the controllable feedforward propagation of axonal transmission in *in vitro* systems (Nguyen et al., 2021; Purwidyantri and Prabowo, 2023; Winter-Hjelm et al., 2023; Winter-Hjelm et al., 2024). A stable, positive fluidic diode effect predicts facilitated nutrient support and growth of cortical projections toward the striatal target, while preventing retro-projections from striatal axons. In addition, the Tesla valve integrates two elegant geometric mechanisms to trap and re-route axons, known as the critical angle (Arrow) effect and the edge guidance effect (Arch), respectively (Renault et al., 2016) (**Figure 1d**). Interestingly, these two mechanisms have been independently proven to facilitate axonal directionality by trapping (Holloway et al., 2019) or re-routing growing neurites (Renault et al., 2016). We therefore reasoned that incorporating fluidic diodicity alongside two geometric constraints, utilizing Tesla valve-shaped microchannels, would ensure the formation of oriented, target-specific corticostriatal projections.

We first investigated whether the Tesla Valve design offers a significant advantage over Arrow and Arch geometries to reconstruct oriented neuronal networks *in vitro*. We produced microdevices with the three designs (Tesla valve, Arrow, Arch; **Figure 2a**) and characterized their fluidic diodicity by tracking the flow of nanobeads within the microchannels after applying various volume differentials (ΔV) between the emitting and receiving chambers (**Figure 2b**). We termed Forward (Fwd) the direct polarization of the fluidic diode (*i.e.*, nanobeads flow from Emitting-to-Receiving chamber); by opposition, we termed Reverse (Rev) the indirect polarization of the diode (*i.e.,* Receiving-to-Emitting). We estimated the diodic effect by calculating the Net velocity of nanobeads (Δv) (see Materials and methods). The Tesla valve configuration showed a stable and positive diodic effect across all ΔV values (F_1,84_=55.33; *p*<0.0001; n=8 microchambers with approximatively 300 moving nanobeads; **Figure 2b,c**). This effect was evident even for the smallest ΔV values, indicating robust performance across all tested volume variations, with mean velocities ranging from +8±2 to +18±2 μm/s. We next compared the fluidic performance of the Tesla valve design with Arrow (critical angle only) and Arch (edge guidance only) geometries. While Arrow microchannels demonstrated an overall positive diode effect, the flow of nanobeads was relatively low, appearing only at high ΔV values (above 15 µL) and remaining inconsistent across higher volumes (**Figure 2b,c**). Fluidic polarization of the Arch design was also inconsistent across ΔV (**Figure 2b,c**). At low ΔV (below 25µL), the flow was even opposite to the expected fluidic diode effect for this configuration, with a maximum velocity reaching -11±1 μm/s at the lowest ΔV (design effect : F_2,30_=27.4, *p*<0.0001; design x volume interaction: F_10,110_=1.9, *p*<0.05; n=8 microchambers).

**Figure 2.**
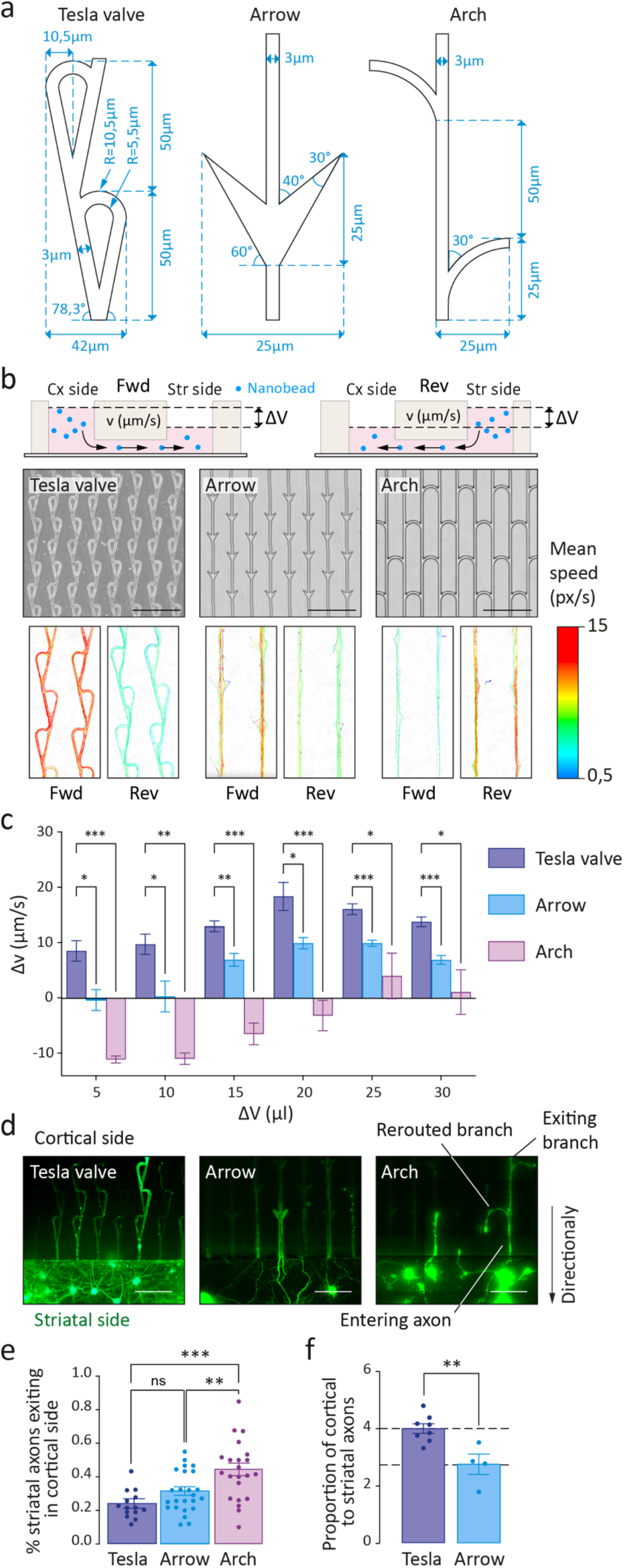
Fluidic and axonal diodicity of Tesla valve, Arrow and Arch geometric designs. **(a)** Schematic of the three geometric designs with dimensional parameters (blue). **(b)** Fluidic diodicity of the three designs using nanobeads flow. ΔV represents the volume differences between cortical and striatal chambers and v the net velocity of nanobeads through microchannels. A Forward (Fwd) flow corresponds to a movement of nanobeads from cortical to striatal chamber. A Reverse (Rev) flow corresponds to the opposite movement. Photomicrographs of the three microchannel designs and their respective contrast phase images showing nanobeads color-coded mean velocity. Scale bar, 150 μm. **(c)** Fluidic diodicity is calculated as the median velocity of nanobeads moving in the forward direction subtracted by the median velocity of nanobeads moving in the opposite reverse direction. A positive value indicates a preferential forward flow for a given ΔV, while a negative value indicates reverse flow (design effect : F_2,30_=27.4, *p*<0.0001; n=8 microchambers; \**p*<0.05, \*\**p*<0.01, \*\*\**p*<0.001). **(d)** GFP-expressing striatal axons back-projecting into the three microchannel designs. Scale bar, 100 μm. **(e)** Proportion of back-projecting GFP-expressing striatal axons toward the cortical chamber (design effect : F_2,57_=9.8, *p*=0.0002; n=20 microchambers; \*\**p*<0.01, \*\*\**p*<0.001; ns, non-significant). **(f)** Proportion of cortical to striatal axons defining global axonal diodicity (*t*-test, \*\**p*<0.01; n=6 microchambers).

We next examined the axonal diodicity potential of the Tesla valve design by evaluating the proportion of cortical axons exiting into the striatal chamber compared with the proportion of striatal axons extending back in the opposite direction (**Figure 2d**).To do so, we expressed the fluorescent protein GFP in either cortical or striatal neurons and reported the number of axons entering and exiting each microchannel, in a fully established and mature network (18 days *in vitro* (DIV18)) (Moutaux et al., 2018b). We found that the Tesla valve design effectively minimized aberrant striatum-to-cortex axonal projections, achieving a 1:5 ratio (1 striatal axon exiting into the cortical chamber for every for 5 axons entering the microchannel; n=14 microfluidic chips from 4 distinct cultures; **Figure 2e**). In contrast, Arrow and Arch geometries only limited the backward growth of striatal axons to a 1:3 and 1:2 ratio, respectively (n=22-24 microfluidic chips from 4 distinct cultures; **Figure 2e**). We further compared the number of cortical and striatal axons reaching the opposite chamber for each channel design (**Figure 2f**). We found that the Tesla valve facilitates the passage of four times more cortical axons compared to striatal axons through the microchannels. In contrast, Arrow microchannels demonstrated significantly lower efficiency, being only half as effective as the Tesla valve geometry in promoting cortical axon passage (p< 0.01; unpaired t-test; n=4-8 microchips from 3 cultures; **Figure 2f**). Our findings demonstrate the efficiency of the Tesla valve in maintaining the intended directional connectivity. This design ensures a reliable diode effect, regardless of volume variations, and features a unidirectional geometric configuration that effectively traps most striatal backward projections while favoring the growth of feedforward cortical axons

### Structural and functional segregation of cortical terminal fields in the striatal chamber

We next investigated the minimum distance required between the two cortical emitting chambers to guarantee distinct separation of their terminal fields in the striatum. In this configuration, a central non-overlapping region where striatal neurons do not receive axonal projections from either cortical compartment should be maintained. This spacing is essential in the brain for preserving distinct functional zones within the striatum and avoiding interference between neighboring pathways (Averbeck et al., 2014; Oh et al., 2014).We estimated the minimum distance between adjacent non overlapping corticostriatal terminal fields to be at least 500 µm, based on previous tracing studies of the rodent corticostriatal system (Mailly et al., 2013; Oh et al., 2014). We therefore designed three devices with non-overlapping zones of 500, 1000 and 2000 µm in length (**Figure 3**). We infected cortical neurons with a lentivirus expressing GFP and quantified axonal fluorescence in the striatal chamber at DIV 18 when the network is already stable and functionally mature [19] (**Figure 3a**). In the 500 µm layout, cortical axons exhibited extensive invasion into the striatal chamber, occasionally extending across to the contralateral corticostriatal terminal field. This pervasive axonal spread resulted in the complete loss of the non-overlapping zone between the two regions. (**Figure 3b**). In contrast, both 1000 µm and 2000 µm configurations efficiently created a zone devoid of axonal projections, as shown by the sudden drop of fluorescence measured in this region (**Figure 3c,d**). Interestingly, the well-defined boundaries of the fluorescent signal indicate that cortical axons do not extend significantly beyond their direct cluster zone and are maintained within a topographically-restricted organization.

**Figure 3.**
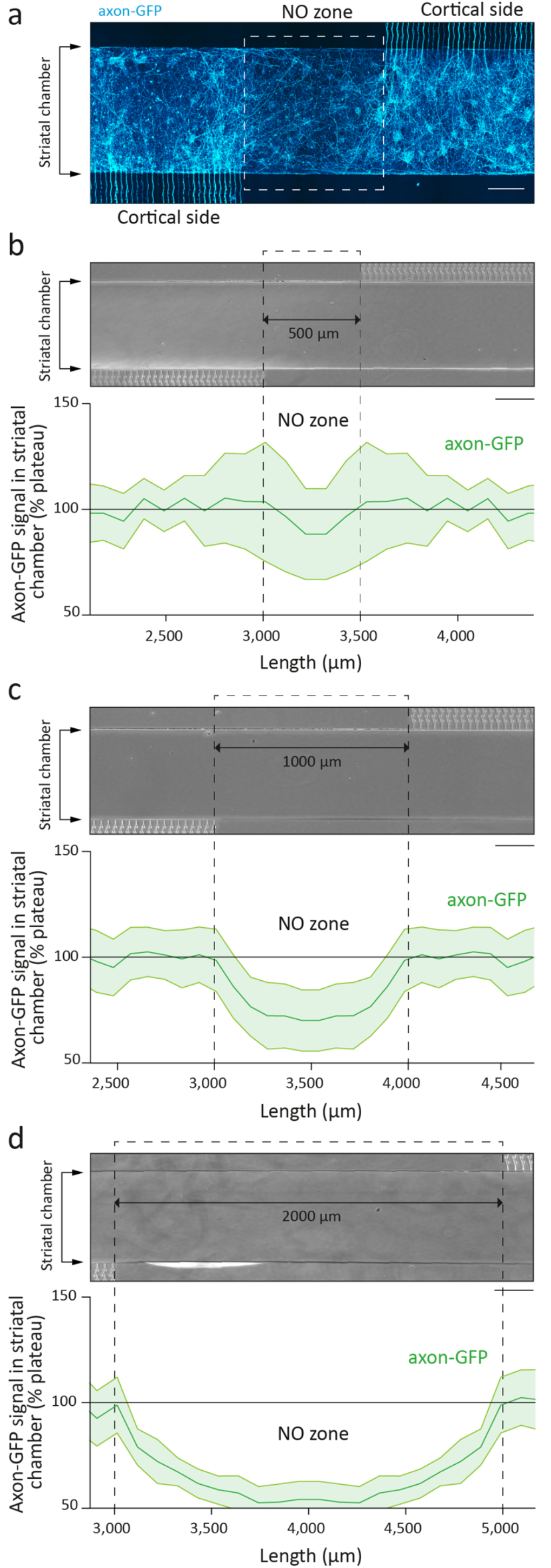
Structural segregation of cortical projections in the striatal chamber. **(a)** GFP-expressing cortical axons (cyan) massively project into the striatal subregion directly facing microchannels. A non-overlapping (NO) zone devoid of cortical projections is created by varying the length of the intermediate zone between microchannels. Scale bar, 250 μm. **(b-d)** Photomicrographs of striatal chambers with varying intermediate NO zone length (**b**, 500 μm; **c**, 1000 μm; **d,** 2000 μm). The efficiency of structural segregation was assessed by plotting fluorescence profiles of GFP-labeled cortical axons centered on the non-overlapping zone for each length. Fluorescence values were normalized to the mean fluorescence in the direct projection zone (n=4 microchambers). Scale bar, 200 μm.

The apparent structural regionalization of the two corticostriatal does not inherently imply functional segregation regarding information transmission and integration. To verify this, we infected striatal neurons with a lentivirus expressing GcaMP6f, a fast, genetically-encoded calcium sensor that monitors neuronal activity (Chen et al., 2013). In culture, striatal neurons predominantly exhibit inhibitory characteristics, showing transient GCaMP6f fluorescence rises only in response to excitatory inputs from cortical neurons establishing functional synaptic connections (Moutaux et al., 2018a; Moutaux et al., 2018b). We took advantage of the fluidic isolation of the microchamber to selectively inhibit neuronal activity of projecting cortical neurons with tetrodotoxin (TTX, a sodium channel blocker) while assessing network activity in the corresponding striatal target region. Calcium imaging was performed in the ventrolateral terminal field region of the striatal chamber (VL St) following selective inhibition of cortical activity, first in the corresponding ventrolateral cortical region (VL Cx) and then in the opposing dorsomedial cortical region (DM Cx) (**Figure 4a,b**). Since VL St neurons primarily receive projections from VL Cx, we reasoned that any residual activity observed after VL Cx inhibition would result from indirect axonal projections from DM Cx (**Figure 4b**). In basal condition, we found that 80±10% of striatal cells displayed GCaMP6f activity in the recorded sub striatal field (500 μm, 79±7%, n=6; 1000 μm, 82±4%, n=16; 2000 μm, 68±2%, n=7), with a mean of 3.5±0.5 events per min per cells for all designs (500 μm, 3.3±0.1%; 1000 μm, 4.1±0.9%; 2000 μm, 3.2±0.1%; **Figure 4c-e**). In the 500 μm configuration, inhibiting the direct projecting VL Cx region is not sufficient to completely abolish GCaMP6f activity in VL St, as 39±11% cells still remained active in the field (F_2,56_=307.4, *p*<0.0001, n=6 microchambers; *p*<0.05 compared to basal), each cell eliciting 0.8±0.2 events per minute in average (F_2,52_=116.8, *p*<0.0001; *p*<0.05 compared to basal; **Figure 4d,e**). However, further addition of TTX in the opposing DM Cx chamber resulted in the complete loss of GCaMP activity in the VL St (percentage of responding cells, *p*<0.001 compared to basal; number of events, *p*<0.001 compared to basal), confirming that the striatal sub region of the 500 μm design receives indirect axonal projections from the opposing cortical chamber. In contrast, both 1000 and 2000 μm configurations showed efficient functional regionalization, as direct inhibition of VL Cx was sufficient to reduce the proportion of responding cells to respectively 2±1% and 3±2% (1000 μm, *p*<0.0001 compared to basal, n=16 microchambers; 2000 μm, *p*<0.0001 compared to basal, n=9 microchambers), with a mean of 0.1±0.05 and 0.1±0.1%, events per cell (1000 μm, *p*<0.0001 compared to basal; 2000 μm, *p*<0.0001 compared to basal; **Figure 4d,e**).

**Figure 4.**
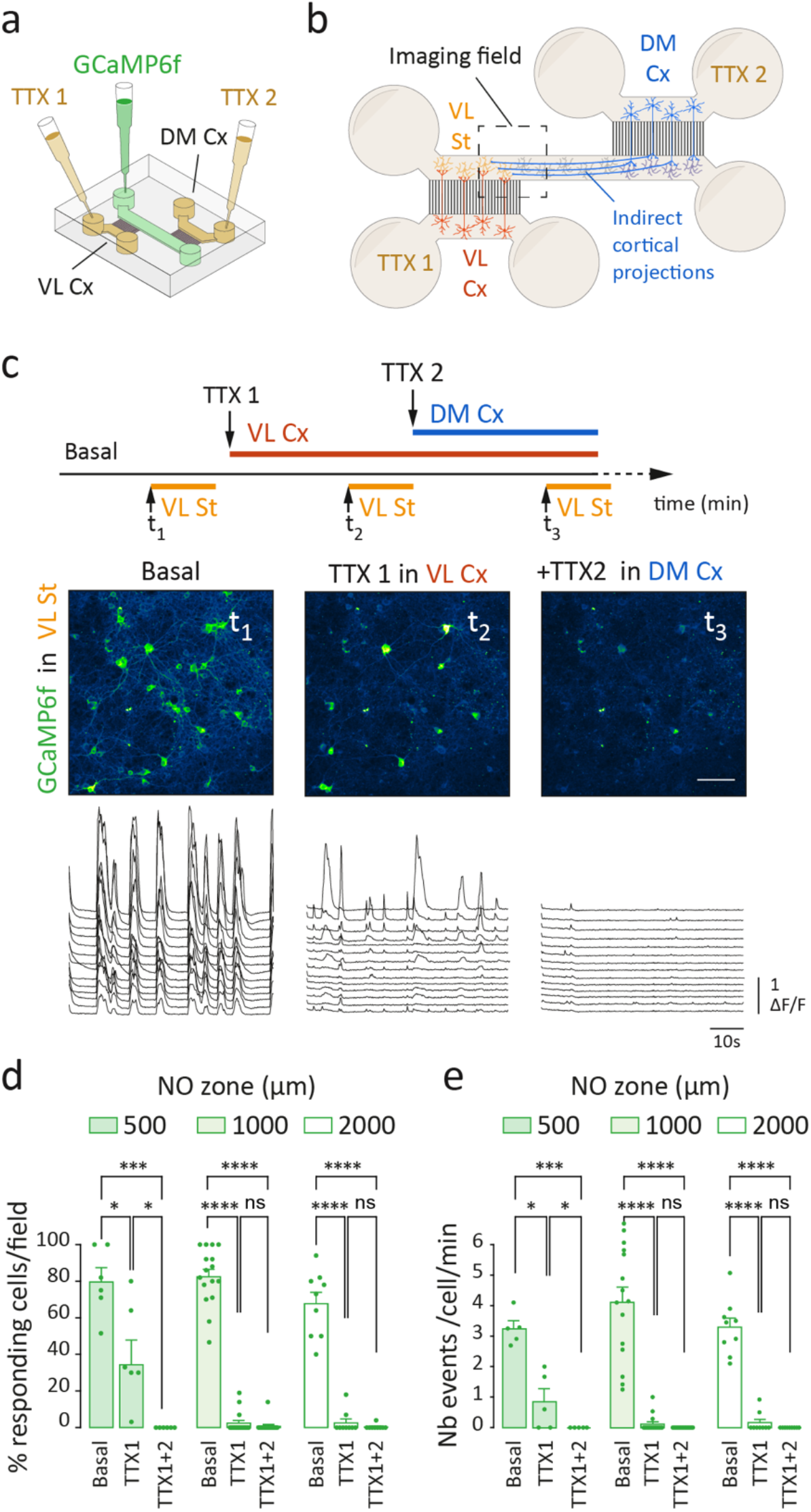
Functional segregation of corticostriatal projections. **(a)** Schematic of the microchip showing sequential TTX injections in the VL Cx compartment (TTX1) and in the DM Cx compartment (TTX2). (b) Striatal GCaMP6f activity was recorded in the VL St zone adjacent to the NO zone to assess the presence of functional connections by DM Cx axons. **(c)** Timeline of sequential TTX injections and GCaMP6f recordings. Microchips were left resting for at least 10 minutes before recording GCaMP6f activity in the VL St at t_1_. TTX was injected in the corresponding VL Cx (5 min) before GCaMP6f recording in the VL St (t_2_). TTX was then injected in the opposing DM Cx chamber before recording in the VL St (t_3_). Live images of GCaMP6f-expressing striatal neurons with representative calcium traces before and after TTX treatments in the 500 μm configuration. Scale bar, 100 μm. **(d,e)** Quantification of striatal GCaMP6f activity responding to cortical inputs for each NO zone configuration, before and after sequential inhibition of cortical activity (TTX1, direct VL Cx inhibition; TTX2, indirect DM Cx inhibition), showed as percentage of responding cells (**d**, length effect : F_2,28_=7.1, *p*<0.005; TTX effect : F_2,56_=307.4, *p*<0.0001; \**p*<0.05, \*\*\**p*<0.001, \*\*\*\**p*<0.0001) and number of events per cell per min (**e**, TTX effect : F_2,52_=116.8, *p*<0.0001; \**p*<0.05, \*\*\**p*<0.001, \*\*\*\**p*<0.0001; ; ns, non-significant; n=6-16 microchambers).

Although both 1000 and 2000 μm configurations efficiently created a non-overlapping zone at both structural and functional levels, the 2000 μm design extended the total length of the striatal chamber to 8000 μm. This dimension proved unsuitable for generating reproducible and homogeneous striatal cultures, as it frequently resulted in uncontrolled flow during cell seeding and medium changes, leading to higher shear stress and disruption of the uniformity of the striatal neuronal culture. Consequently, we selected the 1000 μm configuration with Tesla valve-shaped axonal microchannels to establish a four-node neural network model, in which two parallel corticostriatal pathways topographically project to their respective sub-striatal populations.

### Neurodevelopmental hyperactivity induces profound topographical reorganization of corticostriatal circuits

Several studies have underscored the central role of neuronal activity in the formation and alteration of neural circuits architecture. This is particularly the case within cortical and striatal networks, in which neuronal excitability regulates axonal growth and refining, as well as synaptic connectivity (Katz and Shatz, 1996; Hua et al., 2005; De Marco Garcia et al., 2011). Abnormal cortical activity level, such as hyperexcitability, is a common pathophysiological feature of various neuropsychiatric and neurodegenerative disorders. Alterations in potassium ion channel activity and expression is thought to contribute to the etiopathology of various psychiatric disorders. In particular, genetic mutations of the Kir2.1 potassium channel has been associated with neural network hyperactivity and dysfunctions in bipolar disorder, schizophrenia and autism spectrum disorder (Sicca et al., 2011; Imbrici et al., 2013; Ambrosini et al., 2014; Binda et al., 2018). We therefore investigated the impact of mutant Kir2.1-induced cortical hyperactivity on the global wiring diagram of our topographically segregated corticostriatal pathways. The fluidic isolation of the pre- and postsynaptic compartments provided the opportunity to selectively manipulate cortical neurons within the reconstituted network. We transfected cortical neurons with mutant Kir2.1, in which the GYG motif of the pore region is replaced with three Alanine (Kir2.1^AAA^) (Preisig-Muller et al., 2002). Calcium imaging in the cortical chamber confirmed hyperactivity of Kir2.1^AAA^-expressing neurons, as shown by the 3-fold increase in the number of elicited calcium transients (F_2,42_=40.4, *p*<0.0001; \*\*\*\**p*<0.0001 compared to control; n=15 neurons from 2 microchambers) and the 3.5-fold increase in amplitude after Kir2.1^AAA^ expression (F_2,42_=58.5, *p*<0.0001; \*\*\*\**p*<0.0001 compared to control; **Figure 5a-c**).

**Figure 5.**
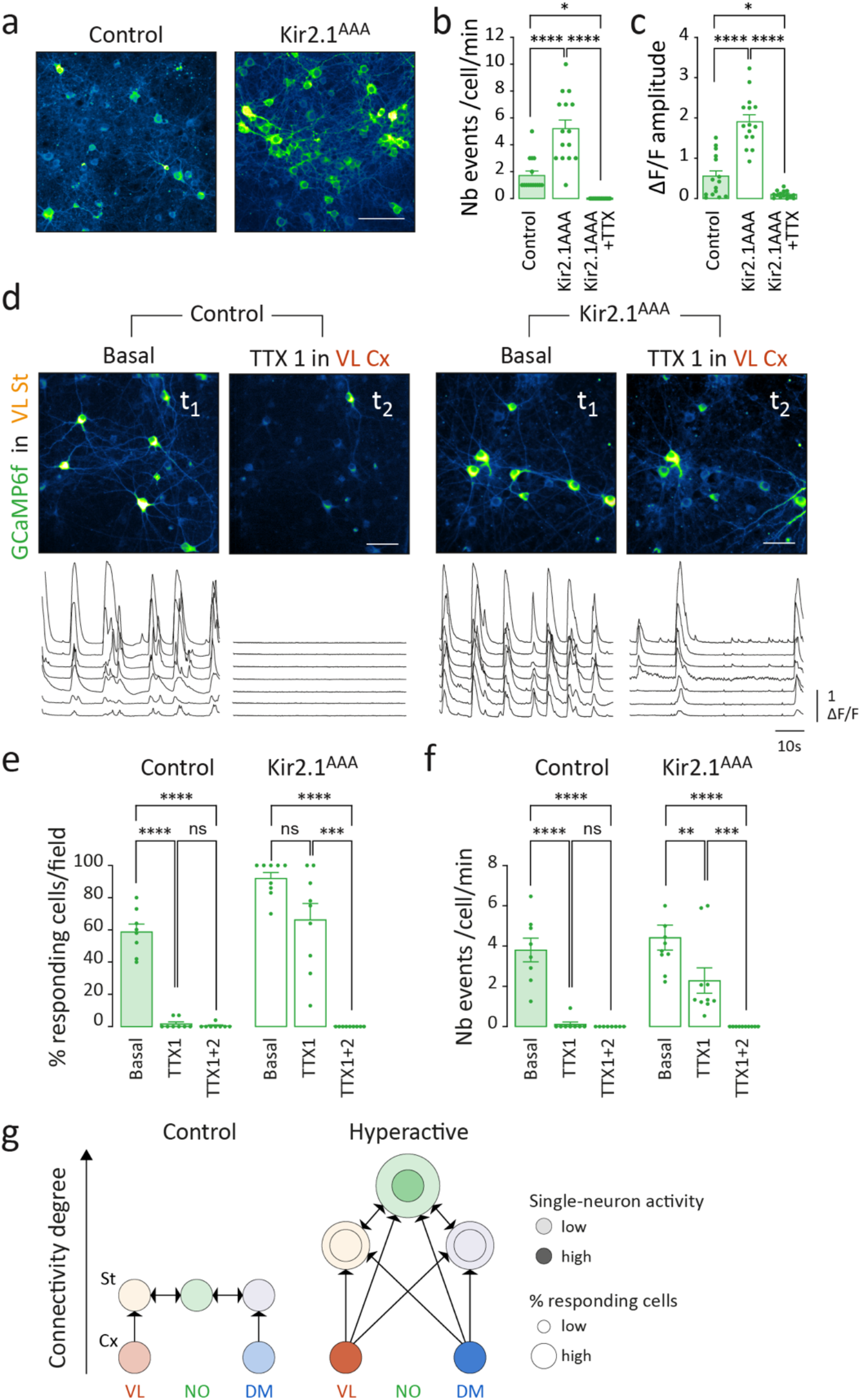
Functional reorganization of hyperactive corticostriatal circuits. **(a)** Live cell images of GCaMP6f-expressing cortical neurons in control and hyperactivated (Kir2.1^AAA^) states. Scale bar, 100 μm. **(b,c)** Quantification of GCaMP6f activity in control and Kir2.1^AAA^-expressing cortical neurons, showed as number of events per cell per min (**b**, F_2,42_=40.4, *p*<0.0001; \**p*<0.05, \*\*\*\**p*<0.0001) and event amplitude (**c**, F_2,42_=58.5, *p*<0.0001; \**p*<0.05, \*\*\*\**p*<0.0001; n=15 neurons from 2 microchambers). **(d)** Live cell images of GCaMP6f-expressing VL St neurons in control and hyperactivated (Kir2.1^AAA^) states before and after TTX treatments in direct VL Cx. Scale bar, 100 μm. Representative calcium traces are shown below. **(e,f)** Quantification of striatal GCaMP6f activity responding to control and Kir2.1^AAA^ cortical inputs, before and after sequential cortical inhibition, showed as percentage of responding cells (**e**, Kir2.1^AAA^ effect: F_1,15_=63, *p*<0.0001; \*\*\**p*<0.0001) and number of events per cell per min (**f**, Kir2.1^AAA^ effect: F_1,15_=4.7, *p*<0.05; \**p*<0.05, \*\*\**p*<0.0001; ns, non-significant; n=8 control and 10 Kir2.1^AAA^ microchambers). **(g)** Theoretical graph representing the topographical organization and connectivity degree between each node of the reconstructed corticostriatal networks, in control and hyperactive conditions. Nodes are interconnected through directional edges. Each node is connected Color intensity of nodes corresponds to single-neuron activity (*i.e.* number of events), while node’s diameter represents the percentage of responding cells in the field. Note the loss of hierarchical organization of hyperactive networks, showing selective changes in global versus individual excitatory drive connectivity reorganization and the emergence of a high-degree convergence node.

The increase in cortical activity has a direct impact on striatal neurons within the field of direct projections, as evidenced by the percentage of responding striatal cells, which rises from 70±8% under control conditions to 92±3% in the Kir2.1^AAA^ condition (F_1,15_=63, *p*<0.0001; \*\*\*\**p*<0.0001 compared to control; n=8-10 microchambers; **Figure 5d-f**). However, our analysis revealed that neither the frequency of events per neuron nor the amplitude of individual responses were significantly altered in the Kir2.1^AAA^ condition compared to the control (3.8±0.6 event/cell/min in control chambers vs 4.4±0.6 event/cell/min in Kir^AAA^ chambers). As expected in the 1000 μm control configuration, the application of TTX to the VL Cx chamber, which harbors direct-projecting neurons, completely abolished GCaMP6f activity in the corresponding VL St neurons (percentage responding cells, \*\*\*\**p*<0.0001 compared to basal; number of events, \*\*\*\**p*<0.0001 compared to basal; n=8 microchambers; **Figure 5e-f**). However, under Kir2.1^AAA^ conditions, significant neuronal activity remained observable in the VL St neurons even after the inhibition of direct cortical projections (basal, 92±3%; TTX1, 66±10% responding cells; n.s). This residual activity cannot be attributed to a reduced efficacy of TTX on hyperactive neurons expressing Kir2.1^AAA^, as TTX completely abolished the activity of Kir2.1^AAA^-expressing neurons in the cortical chamber (\*\*\*\**p*<0.0001 compared to Kir2.1^AAA^; **Figure 5b,c**). When TTX was applied to the opposite DM Cx chamber, GCaMP6f activity was entirely eliminated in the VL St (\*\*\*\**p*<0.0001 compared to basal; \*\*\**p*<0.001 compared to TTX1), confirming that adjacent axonal projections invaded the target zone.

We next explored the functional disorganization resulting from neurodevelopmental hyperactivity by building a graph theoretical analysis based on the GCaMP6f data (**Figure 5g**). We found that cortical hyperactivity primarily leads to the recruitment of additional striatal neurons within both the projection and the non-overlapping zone, rather than increasing the number of synaptic events or the strength of responses in individual neurons. This redistribution of connectivity results in a greater excitatory drive at the global population level, while maintaining similar activity profiles at the single-neuron level. Axonal invasion of the non-overlapping zone further destabilizes the topographical organization of the network, creating dysfunctional convergence nodes that receive inputs from opposing cortical regions. The recruitment of additional cells in both the target and the adjacent striatal region, without significantly changing the intrinsic activity of each neuron, may therefore enhance and expand network responsiveness, disorganizing the integration and processivity of information propagating within the network, without altering single-neuron properties (**Figure 5g**).

We next investigated when and how functional reorganization occurs in hyperactive networks. The standardized architecture of microchips allows to perform detailed, longitudinal analysis, tracking network development from early formation through to late maturation stages. We analyzed the structural dynamics of axonal branching and connectivity thanks to the spatiotemporal compartmentalization of the co-culture. We focused on key developmental time points [4, 7, 10, 14, and 21 days in vitro (DIV)] representing critical stages of network formation and maturation in microchips (Moutaux et al., 2018b). We mapped the structural evolution of corticostriatal projections using tdTomato, a fluorescent marker co-expressed with Kir2.1^AAA^, which is cleaved off to diffuse freely throughout the cytoplasm, including into axonal compartments. Microchips were systematically subjected to high-resolution microscopy scanning of the whole striatal chamber to track and quantify structural evolution of the network overtime (**Figure 6a**). We observed that by DIV4, axonal projections from Kir2.1AAA-expressing neurons had already reached the striatal target zone, while control axons were still in the process of crossing the microchannels. In the control group, axonal fluorescence continued to increase steadily within the striatal target zone directly facing the microchannels between DIV7 and DIV14. From DIV18 onward, the control network began to stabilize and mature, corroborating our previous findings with reconstructed corticostriatal circuits on microchips (Moutaux et al., 2018b; Virlogeux et al., 2018). After DIV21, no further increase in axonal signal was observed within the striatal chamber, indicating that the network had reached structural maturity. At this stage, the network maintained a distinct topographical organization, with the majority of axonal fluorescence localized to the direct projection zone. In contrast, Kir2.1^AAA^-expressing microchambers showed a continued increase in axonal signal across the entire striatal compartment, with significant invasion of the non-overlapping zone beginning at DIV14. By DIV18, the topographical organization of these hyperactive networks had fully collapsed, leading to the formation of a homogeneous projection zone that spanned the entire striatal compartment, including the non-overlapping region. This reorganization resulted in the creation of an intercortical convergence node, receiving inputs from both opposing cortical regions and disrupting the original segregation of projections.

**Figure 6.**
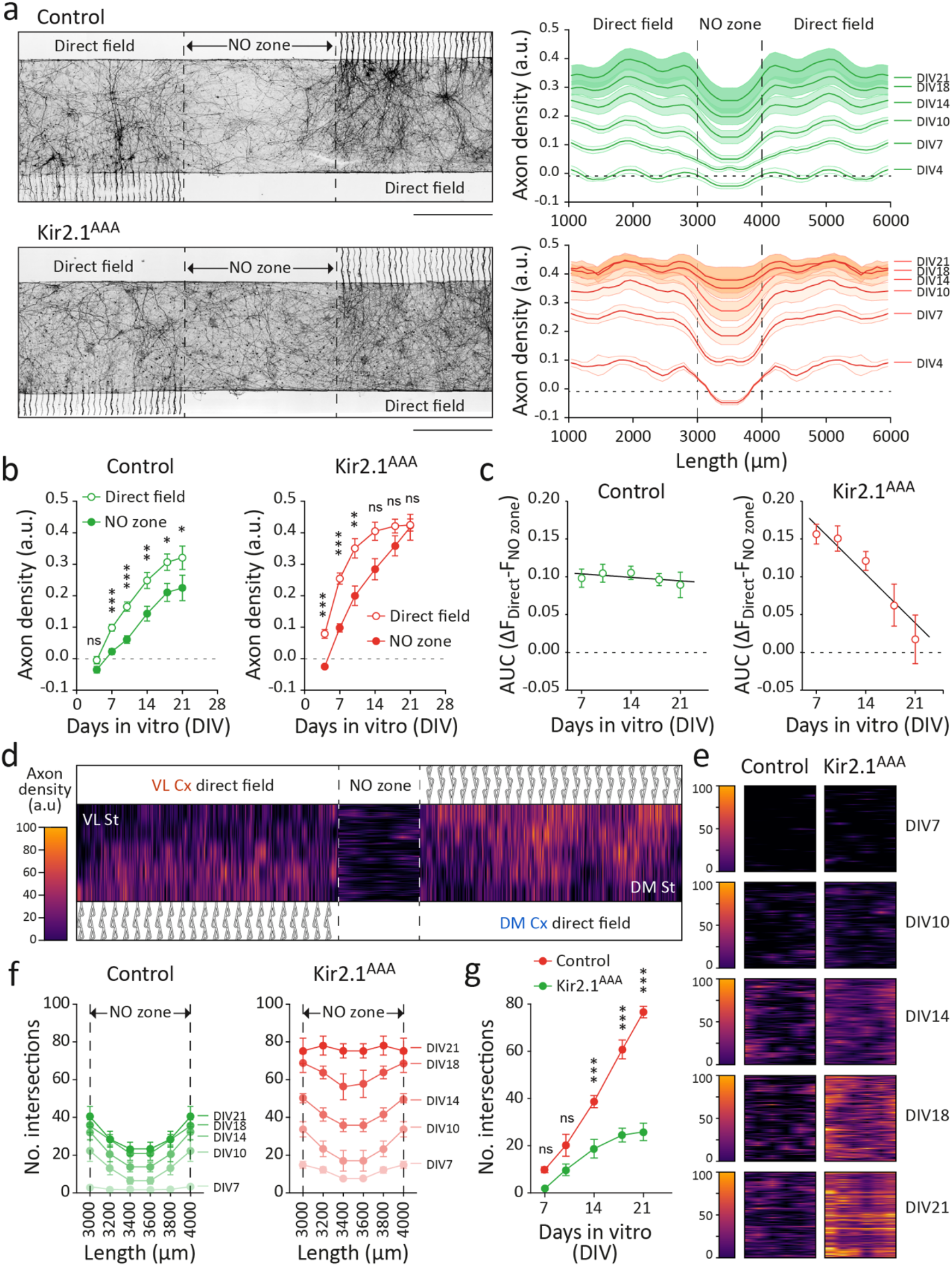
Spatiotemporal reorganization of hyperactive corticostriatal networks during development. **(a)** Confocal images of control and Kir2.1^AAA^-expressing cortical axons projecting into the striatal chamber. Note the axonal invasion of the NO zone by Kir2.1^AAA^ cortical axons. Scale bar, 500 μm. Fluorescence profiles of cortical axons centered on the non-overlapping zone are plotted for each developmental time point (4, 7, 10, 14, 18 and 21 DIV; n=16 microchambers per DIV). **(b)** Average axon fluorescence in the direct projecting fields and in the NO zone during network development obtained from fluorescence profiles in **(a)**. Control conditions show similar growth kinetics between the direct projection field and the NO zone (F_5,147_=0.48, ns), despite constant stronger signal in the direct projecting field (F_1,52_=28.7, *p*<0.0001; \**p*<0.05, \*\**p*<0.01, \*\*\**p*<0.001). Altered growth kinetics in Kir2.1^AAA^-epxressing axons between the direct projecting fields and the NO zone (F_5,129_=2.42, *p*<0.05; \*\**p*<0.01, \*\*\**p*<0.001, ns. non-significant). **(c)** Area under the curves from **(b)** were used to calculate signal amplitudes between the direct and NO zones in control and Kir2.1^AAA^ networks. Note that a negative slope is representative of a constant increase in axon invasion in the NO zone. **(d)** Representative heatmap of cortical axon branches in direct projecting fields and in the NO zone of the striatal compartment (see Materials and Methods for details). **(e)** Heatmap representations of the NO zone in control and Kir2.1^AAA^ networks for each developmental time point. **(f)** Linear Sholl analysis of the NO zone showing the rapid increase in the number of intersections, in Kir2.1^AAA^ conditions. **(g)** Average number of intersections in the NO zone at each developmental point showing accelerated axonal invasion by hyperactive cortical axons (F_4,75_=18.6, *p*<0.0001; \*\*\**p*<0.001; ns, non-significant; n=5 microchambers).

Interestingly, we observed that the kinetics of axonal invasion between the direct projection field and the non-overlapping zone were significantly altered by cortical expression of Kir2.1^AAA^. In the control group, axonal fluorescence increased similarly in both regions, though the signal consistently remained stronger in the direct projection zone, maintaining the segregated topographical organization of parallel corticostriatal circuits throughout network formation and maturation (**Figure 6b**). This created a stable difference in signal amplitude between the direct and non-overlapping zones, which was preserved over corticostriatal networks formation and maturation (**Figure 6c**). Axons in the Kir2.1^AAA^ condition followed a similar growth trajectory within the direct projection zone, although their density was both amplified and accelerated. By DIV14, the axonal signal in the direct projection zone had already reached the maximum level observed in the control group by DIV21, stabilizing thereafter, much like in the control condition. However, axonal fluorescence in the non-overlapping zone increased linearly from DIV7 on, reaching its peak at DIV21 (**Figure 6b**). This linear increase in density suggest that neuronal hyperactivity not only accelerates axonal growth from the earliest stages of development, but also drives continued expansion, eventually leading to the invasion of the non-overlapping zone, equivalently saturating all neurons within the striatal chamber with connections (**Figure 6c**).

We next utilized the space-time compartmentalization of neuronal networks on a chip to implement a custom linear Sholl analysis method, designed to capture the dynamics of axonal growth within the non-overlapping zone (**Figure 6d**). This method offers the advantage of providing a reproducible, quantitative, and automated longitudinal analysis of topographical network reorganization in response to genetic perturbations. To achieve this, we systematically segmented the microchambers with horizontal lines extending from the end of the microchannels to the opposite wall for the two direct projection zones, and with vertical lines across the width of the non-overlapping zone (see Materials and methods). Each axonal branch crossing these defined lines was recorded and integrated into a matrix mapping the entire striatal chamber (see Materials and methods for custom code). This heatmap representation of axonal density enabled rapid comparison of data for each neuronal compartment at defined developmental time points (DIV7, 10, 14, 18, and 21; **Figure 6e**). To better understand the reorganization kinetics of corticostriatal circuits, we focused specifically on the non-overlapping zone. The analysis revealed a striking effect of Kir2.1^AAA^ expression on axonal growth and branching dynamics outside the direct projection zone. While the non-overlapping zone remained relatively devoid of axonal branches in control conditions—particularly in its central region—hyperactive Kir2.1^AAA^-expressing axons began invading the entire area as early as DIV7, expanding from the periphery toward the center. Unlike the control group, where axonal growth rapidly reached a stable state, hyperactive networks continued expanding until a peak was reached at DIV21. By this point, the central non-overlapping zone contained 3.2 times more Kir2.1^AAA^ axons than in controls (75±4 axons in the Kir2.1^AAA^ group versus 23±3 axons in the control group, p<0.0001, *t*-Test; **Figure 6f**). This dramatic increase in axonal density was primarily driven by accelerated growth dynamics, with hyperexcitable axons growing at a rate 2.6 times faster than in control conditions (∼1.8 new axons/day in the control group versus ∼4.7 new axons/day in the Kir2.1^AAA^ group; **Figure 6g**). These observations suggest that under conditions of hyperexcitability, corticostriatal networks not only fully invade their target regions but also rapidly expand into adjacent zones, eventually occupying the entire 7000 μm striatal compartment. This accelerated and extensive axonal growth leads to a loss of topographical organization, contributing to the structural and functional disorganization of network activity.

## DISCUSSION

Studying the complex reorganization of topographically wired neural networks remains a significant challenge, yet it is crucial to advancing our understanding of neurodevelopmental diseases. The primary difficulty lies in developing physiological models that preserve this intricate architecture, while providing access to external and/or genetic perturbations. Here, we present a biologically grounded model of neuronal network using a microfluidic on-a-chip device suitable for the longitudinal analysis of neurodevelopmental perturbations. By applying this device to hyperactive corticostriatal networks, we identified complex local and long-range network reorganization, highlighting early topographical reorganization and the emergence of dysfunctional convergent nodes.

### Reorganization of the wiring diagram in hyperactive networks creates new convergent nodes

Hyperactivity and disorganization of corticostriatal circuits are key features of various neurodevelopmental and psychiatric disorders (Shepherd, 2013; Wolfram and Baines, 2013; Nelson and Kreitzer, 2014). Despite their importance, understanding how, when and where these progressive defects occur remains elusive due to the technological limitations of *in vivo* studies. By transposing the topographical organization of corticostriatal networks onto a chip, we identified that accelerated growth of cortical projections withing striatal territories is a primary driver of the loss of topographic organization within hyperactive networks, resulting in the formation of dysfunctional convergence nodes.

Remarkably, although hyperexcitable axons increased the local density of connections at the population level in the target zone, this reorganization did not result in over-connection at the single-neuron level. This selective effect may be attributed to local competition between neighboring hyperexcitable axons, preventing over-connection on individual neurons (Hua et al., 2005; Singh and Miller, 2005). Instead, local connections redistributed across a broader target area, expanding excitatory input onto striatal territories without altering the electrophysiological properties at the single-neuron level. In addition to disrupting the local topography, cortical hyperexcitability also prompted the recruitment of neurons in adjacent regions. This invasion into previously unconnected territories expanded the network’s connectivity map well beyond its original boundaries, significantly altering the balance between local and long-range connectivity. These findings align with prior studies where neuronal activity inhibition via Kir2.1 overexpression in cortical neurons reduced axonal growth and branching in contralateral target zones. However, in that case, hypoexcitable axons extended beyond their target area without forming stable connections (Wang et al., 2007). This distinction may arise from the differences in cortical networks investigated. In corticocortical circuits, the balance between pre- and post-synaptic activity is crucial for the proper formation of axonal projections (Huang et al., 2013; Suarez et al., 2014). In corticostriatal circuits, however, cortical axons connect to inhibitory GABAergic medium spiny neurons, which are highly hyperpolarized and electrically silent, both *in vitro* and *in vivo* (Nisenbaum and Wilson, 1995; Wilson and Kawaguchi, 1996). This disparity in membrane excitability between pre- and post-synaptic neurons in corticostriatal networks may accelerate the growth kinetics of cortical neurons as they invade the striatal compartment, leading to faster and more extensive cortical axonal expansion until all neurons in the adjacent region are connected to projections from neighboring areas.

The extension of hyperexcitable axons into adjacent regions, without altering the properties of individual neurons, suggests that reorganized nodes—though regionally aberrant—could still process information in a physiologically plausible manner. This may lead to the formation of new functional convergence nodes, potentially enabling the processing of information from unrelated cortical areas. Interestingly, axonal extension beyond the original target area has also been observed under physiological conditions, such as during extensive learning. Studies in primates have shown that axonal projections involved in tool-use motor learning can extend over long distances (several millimeters) to invade areas typically devoid of connections in naïve animals (Hihara et al., 2006). In humans too, examples of large-scale axonal remodeling of cortical circuits have been reported. For example, the visual system of subjects trained to juggle undergo significant structural remodeling of the white matter, suggesting selective reconnections in cortical circuits (Draganski et al., 2004; Scholz et al., 2009). Extensive rearrangement of brain circuits after years of intensive training is also associated with the exceptional skills of professional pianists (Bengtsson et al., 2005) and of extreme foot users (Dempsey-Jones et al., 2019). This large-scale rewiring of hyperactive cortical circuits is believed to create new functional modules that support the acquisition of specific motor skills (Johansen-Berg, 2007; Bennett et al., 2018).

### Implications for neurodevelopmental and psychiatric diseases

These findings are also highly relevant for understanding neurodevelopmental connectopathies characterized by hyperactive regions—whether due to external perturbations or genetic mutations affecting neuronal excitability, where hyperactivity in certain cortical and striatal regions can lead to aberrant network expansion. This highlights the necessity for future studies to focus on specific subregions and suggests strategies for targeted therapeutics (Steinberg et al., 2015). For instance, autism spectrum disorders are characterized by a shift in the balance between local and long-range connectivity during embryonic and postnatal development, with hyperconnectivity observed in frontocortical circuits (Supekar et al., 2013; Allen and Morishita, 2024). Genetic risk factors associated with neurodevelopmental disorders also point to an imbalance between local and long-range inputs. For example, mutations in Mef2c, linked to both schizophrenia and autism spectrum disorders, result in excessive callosal long-range excitatory inputs (Rajkovich et al., 2017).

Current psychiatric medications offer only moderate symptom relief and do not effectively alter the course of disorders. Since many of these drugs were developed based on clinical observations, little is known about how they produce effects at the circuit level to alleviate symptoms. Antipsychotic drugs, for example, alleviate only a subset of schizophrenia symptoms, without addressing the primary cause of circuits dysfunctions (Hyman, 2012). Research increasingly suggests that intervention during neurodevelopment may be the most effective way to influence the course of psychiatric disorders (Krol and Feng, 2018). Given the dynamic changes in synapse number, function, and connectivity during development, intervening during these critical periods may help support normal development and provide lasting benefits. Our findings also indicate the need for rethinking current approaches to treatment design. Regionally targeted interventions, rather than broadly reducing network activity through systemic treatments may provide more effective-therapies for conditions marked by hyperactive, abnormal networks (Steinberg et al., 2015). Next-generation therapies, including pharmacological, electrical and optical strategies, should therefore aim at specific circuits, particularly those addressing dysfunctional convergence nodes. However, developing such circuit-based therapeutics requires a deeper understanding of the complex mechanisms governing network connectivity. This gap is further widened by the limited knowledge of how circuit connectivity and function mature in disorders with developmental components. To address these limitations, developing biologically grounded models of complex neural networks that allow for selective manipulation and direct observation of connectivity dynamics throughout development may offer unique platforms for advancing next-generation therapies.

### A minimalistic platform to analyze brain circuits architecture in health and neurodevelopmental diseases

The brain is increasingly recognized as a system of interrelated networks that follow intricate topographical patterns. This shift in understanding highlights the importance of examining brain connectivity from a network-centric perspective, rather than viewing it as separate circuits (Sporns, 2014). Corticostriatal pathways, in particular, demonstrate this complexity through their organization into subnetworks, where specific zones allow for the convergence and divergence of information (Flaherty and Graybiel, 1991; Berendse et al., 1992; Hoffer and Alloway, 2001; Mailly et al., 2013; Averbeck et al., 2014; Hintiryan et al., 2016). However, the complex nature of topographically structured neural networks has long posed a significant challenge for neurodevelopmental disorders. Traditional *in vitro* models have struggled to capture the full complexity of these networks, often falling short in replicating both their structural and functional aspects (Zhao et al., 2020). This limitation has hindered our ability to gain deeper insights into the mechanisms underlying circuit reorganization, a crucial component in understanding various neurodevelopmental conditions. Our platform addresses this longstanding issue by providing a more accurate and dynamic representation of how neuronal circuits respond to genetic or environmental perturbations. One key advantage of microfluidic platforms is the precise control of geometric parameters and environmental factors it offers, allowing for highly controlled experiments and a better simulation of how neuronal circuits respond to genetic and environmental changes. Moreover, monolayered cultures on a single optical plane design enables clearer observations, which improves spatial and temporal imaging capabilities. This feature, along with the integration of advanced analytical tools, facilitates real-time observation of axonal reorganization during development, offering insights into both local and long-range circuit dynamics. Another important feature of microfluidic on-a-chip platforms is the standardized nature of culture conditions, enhancing consistency across experiments, and improving the reliability of findings in neurodevelopmental studies.

This novel approach may serve as a complementary tool to *in vivo* and human imaging studies, offering a minimalistic yet highly informative model of neural network formation and rearrangement. By bridging the gap between oversimplified *in vitro* models and the complexity of *in vivo* systems, our platform opens new avenues for investigating the effects of genetic and environmental factors on neural circuit development, and may help biomedical actors to develop next-generation neurotherapies. Our minimalistic platform may serve as an effective pre-screening tool that can be adapted to human neurons to develop characteristic changes similar to what is observed in neurological disorders before transitioning to more advanced and complex *in vivo* models. This strategy may allow precise mechanistic characterization in a bottom-up efficient approach to study neural network alteration and dysfunctions. By providing a more accurate representation of neuronal circuit response to genetic perturbations, our platform may therefore open new avenues for investigating neurodevelopmental disorders.

## MATERIALS AND METHODS

### Microfluidics fabrication

Polydimethylsiloxane (PDMS) microfluidic devices were fabricated and prepared as previously described (Virlogeux et al., 2018). Briefly, a master mold was made with SU-8 photoresist on a silicon wafer using a dual thickness photolithography process. A first layer of SU-8 2005 (5μm high) was spin-coated on a 3-inch silicon wafer, then baked, exposed and developed for defining the microchannels. A second layer of SU-8 2100 (100μm high) was subsequently spin coated and aligned over the first level and was developed to build seeding chambers. Note that a short silicon over-etching was performed with reactive ion etching between the two steps so that the microchannels pattern was visible for alignment of the second mask. Silicon wafers were then replicated using epoxy resin (resin epoxy R123, SoloPlast) and PDMS microfluidic chips were prepared using a 10:1 mix of silicon elastomer and its curing agents (Sylgard 184, Dow Corning). Air bubbles were removed by vacuum for 1h and PDMS polymerization was achieved at 70 °C for 1 h 30 minutes. PDMS microfluidics chips were then unmolded, cut to create seeding wells, cleaned by sonication in 100% ethanol, rinsed with distilled water and dried at 60 °C. Bonding the chips with FluoroDish (FD35-100, ThermoFisher Scientific) were performed using a low-pressure plasma treatment (Femto plasma cleaner, Diener Electronic, Germany) for 30 seconds for surface activation. Bonding was sealed at 60°C for 15 minutes to create covalent bonds between the microfluidics chips and the FluoroDish. Microfluidic devices were then subjected to a 10 second plasma treatment just before the coating. Coating was performed overnight at 4°C with poly-D-lysine (0.1 mg/ml) in the cortex chambers and with a mix of poly-D-lysine (0.1 mg/ml) and laminin (5 μg/ml) in the striatal chamber. The microchips were carefully washed before plating with growth medium (Neurobabasal medium supplemented with 2% B27, 1% Glutamine 200 mM, and 1% penicillin/streptomycin) to remove excess coating, and were placed at 37°C before neurons were plated. Microchambers were carefully inspected every 2-3 days and before experiments.

### Nanobeads videotracking

Polystyrene beads (diameter 500 nm, cat. #59769, Merck) were diluted at final concentration of 1:5000 (v/v) in MilliQ water. Beads in solution were sonicated for 20 minutes to prevent clusters. Microfluidic chips were activated trough plasma atmosphere at 4 mbar air for 2 minutes. Each chamber was pre-filled with 10 µL of MilliQ water to assure homogenous filling and flow stabilization inside microchannels. Acquisitions were made on a custom inverted microscope (N.A_illumination_=0.3) with a ×60 oil objective (N.A_detection_=1.3, Olympus). Wavefront images were acquired using a custom Quadrivalve Lateral Shearing Interferometry (QLSI) camera system at a 15Hz frame rate (Bon et al., 2009).

### Primary neuronal cultures

Cortex and ganglionic eminences were dissected from e17.5 rat embryos and digested enzymatically with trypsin, followed by two incubations with trypsin inhibitor solutions, and finally dissociated mechanically. Dissociated striatal and cortical neurons were resuspend in growth medium (5 x 10^6^ cells in 140 μl for cortical neurons, and in 70 μl for striatal neurons), and plated in seeding chambers at a final density of 3500 cells/mm^2^ for cortical neurons, and 7000 cells/mm^2^ for striatal neurons. Cells were left in the incubator for at least 3h to allow cell adhesion, then all compartments were carefully filled with growth medium.

### Constructs and viruses

Neurons were either infected with lentiviruses (LV) and adeno-associated viruses (AAV) one day after plating (DIV 1) for 24h, or were electroporated with plasmids using Amaxa Nucleofactor (Lonza) before plating. The following constructs, LVs and AAVs were used for the study: pRRLSIN.cPPT.hPKG-GFP (Addgene #12252),), pCAG-Kir2.1Mut-T2A-tdTomato (Addgene #60644), AAV9.CamKII.GCaMP6f (Addgene #100834-AAV9), pLV.PKG.GFP (Addgene #12252-LV) and pLV.CMV.Cherry (Addgene #36084-LV).

### Immunostaining and confocal imaging

Neurons were fixed by filling the microchambers with PFA/Sucrose (4%/4% in PBS) for 20 min at room temperature (RT) followed by PBS washes and were incubated for 1h at RT with a blocking solution (BSA 1%, normal donkey serum 2%, Triton X-100 0.1%). Cortical and striatal chambers were then incubated with anti-GFP (ThermoFisher Scientific, #A10262, 1:1000) polyclonal primary antibody overnight at 4 °C and appropriate fluorescent secondary antibodies were incubated for 1h at RT. The immunofluorescence was maintained in PBS for a maximum of one week in the dark at 4 °C before confocal acquisitions. Immunostainings were acquired with a ×10 objective (0.3 NA) using an inverted confocal microscope (Eclipse Ti2, Nikon) coupled to a spinning-disk confocal system (CSU-W1, Yokogawa) connected to a wide field sCMOS camera (Orca-Fusion BT, Hammamatsu). GFP-positive axons were imaged in the striatal chamber using multidimensional (x,y,z) acquisitions and were reconstructed as a single mosaic with a 25% (x,y) overlap. For axonal directionality experiments, microchips were maintained at 37 °C and 5% CO2 and live axons expressing endogenous GFP were imaged in microchannels with a ×60 objective (1.49 NA). The system is driven by NIS-Elements (Nikon Instruments)

### Live cell videomicroscopy

Live-cell recordings of GCaMP6f signals were performed using an inverted microscope (Axio Observer, Zeiss) coupled connected to wide field hight sensitivity sCMOS camera (Prime BSI Express, Photometrics) and maintained at 37 °C and 5% CO2. GCaMP6f images were taken every 200 ms for 60s using a ×25 oil-immersion objective (0.8 NA). The system is driven by Metamorph Imaging System Premier (Molecular Devices).

### Quantifications and image analysis

For nanobeads flow analysis, wavefront images were reconstructed using an open-source MATLAB toolbox (available at https://github.com/baffou/PhaseLAB). The flow of beads was detected and tracked using an open-source ImageJ plugin (Ershov et al., 2022). Temporal average values were subtracted to the individual frames to remove static components of the movies and the position of each bead was instantly analyzed on each frame using custom MATLAB scripts. Events that were tracked for less than 20 frames were considered as outliners. We estimated the Net velocity of nanobeads (Δv) by calculating the difference between Forward and Reverse average velocity of nanobeads for each volume differentials (ΔV). At least 8 microchambers were imaged, with approximatively 300 moving nanobeads per acquisitions for each ΔV. For the analysis of axonal directionality, live imaging of GFP-expressing axons was performed at DIV4, 7, 10, 18, and 21. We estimated axonal directionality at DIV18, by quantifying the number of GFP-positive axons entering the microchannels from one compartment and exiting into the other compartment (striatum-to-cortex, or cortex-to-striatum). Axonal directionality was then estimated by calculating the proportion of exiting axons relative to the number of axons entering. Each design was tested using 10 fields per chamber from at least 14 microchambers prepared from 4 independent cultures. Global axonal diodicity was evaluated by calculating the proportion of striatum-to-cortex axons relative to the number of cortex-to-striatum axonal projections from at least 4 microchambers prepared from 3 independent cultures. Live cell calcium analysis was performed between DIV18 and DIV21. Neurons were left to rest for 10 min before recording. Tetrodotoxin (TTX, 5μM) was diluted in fresh medium and applied in the cortical compartment using inter-chamber differential pressure, which limits the diffusion of liquids between the striatal and cortical chambers. Cortical neurons were incubated for at least 5 minutes at 37°C before live cell imaging. Calcium fluorescence images were processed using ImageJ (FIJI v2.1.0, https://imagej.nih.gov/ij/) and analyzed using Matlab 2014b. Regions of interests (*i.e* neuronal soma) were delineated manually and ΔF/F traces were calculated using FluoroSNNAP software for Matlab (Patel et al., 2015). Regions of interest were conserved between acquisitions to follow the response of the network between different conditions. Calcium event detection was performed with homemade functions (available at https://github.com/PoinsotM/Code). Acquisition fields were randomly distributed along the striatal chamber and all responding cells in the field were defined as regions of interest. Events were detected when the ΔF/F value was over five standard deviations of the ΔF/F background signal. The program detected the number and amplitude of each event. Each condition was tested using at least 6 microchambers from 6 independent cultures for evaluating the 500, 1000 and 2000 μm designs. Kir2.1^AAA^ functional effect was analyzed using at least 8 microchambers from 5 independent cultures, except for cortical calcium activity where 2 microchambers prepared from one culture were used. For axonal density analysis in the striatal compartment, mosaic reconstruction was analyzed using ImageJ (FIJI v2.1.0). A region of interest encompassing the entire striatal chamber, centered within the non-overlapping zone, was used to quantify the overall fluorescence signal. After background subtraction, the signal was plotted as a function of the longitudinal coordinates of the chamber. Each design (500, 1000 and 2000 μm) was tested using at least 4 distinct microchambers from 4 independent cultures. Kir2.1^AAA^ structural effect was tested using at least 16 microchambers per DIV and per condition from 9 independent cultures. The linear Sholl analysis was performed by segmenting the striatal compartment with horizontal lines from the end of the microchannels to the opposite wall for the two direct projection zones every 180 μm, and with vertical lines across the width of the non-overlapping zone every 166 µm, using ImageJ (FIJI v2.1.0). Image were binarized and the color scale was inverted so that cortical axons invading the area had a pixel value of 255, while the background was assigned a value of 0. Pixel intensity profiles (0 or 255) were extracted along each line to estimate the number of axonal intersections. These intensity profiles were also used in a custom Python algorithm to provide a heatmap representation of axonal density probability at each DIV (code available at https://github.com/PoinsotM/Code). Five microchambers from 16 microchambers were randomly distributed at each DIV.

### Statistical analyses

Statistical analyses were performed using Prism10 (GraphPad Software). The fluidic diode effect (nanobeads), functional regionalization of the three designs (500, 1000, 2000 μm), functional invasion of Kir2.1^AAA^ in the non-overlapping zone, the kinetics of axonal invasion between the direct projection field and the non-overlapping zone, and the linear Sholl analysis overtime were analyzed using two-way ANOVA followed by a Tukey post hoc test. Axonal unidirectionality studies (Tesla, Arrow, Arch), and cortical calcium activity were analyzed using one-way ANOVA followed by a Tukey post hoc test. Global axonal diodicity between Tesla and Arrow geometries were compared using unpaired two-tailed Student’s t-test. Results are expressed as mean ± SEM. The criterion for statistical significance was set at *p*<0.05.

## ACKNOWLEDGMENTS

We thank Kim Mormentyn, Amelle Nasri and Catherine Lepolard for technical support; Genevieve Rougon for critical comments on the manuscript; Sophie Halliez and members of the lab for helpful discussions; E. Nivet’s team (INP, CNRS, Aix Marseille Université, UMR7051) and the NeuroBioTools cloning facility (INT, CNRS, Aix Marseille Université, UMR7289) for lentivirus production; NCIS imaging facility at INP for image acquisitions; and the Neurotechnology Center at INT for microchip production, and live cell imaging. This work was supported by grants from the Agence Nationale de la Recherche (ANR-19-CE16-0027, ANR-22-CE17-0034), Association Nationale de la Recherche et de la Technologie (ANRT-CIFRE-2021/1043), Fondation France Alzheimer (2021-6239), Institut Carnot STAR, and Fond d’Investissement INT. This work has received support from the French government under the Programme «Investissements d’Avenir», Initiative d’Excellence d’Aix-Marseille Université via AMIDEX funding (AMX-19-IET-004), and ANR (ANR-17-EURE-0029).

## AUTHOR CONTRIBUTIONS

Conceptualization, M.P. and M.C.; Methodology, M.P., B.M., B.C. and M.C.; Investigation, M.P., M.D.S., B.M., A.B.C. and B.C.; Formal Analysis, M.P., B.M., M.D.S. and M.C.; Writing – Original Draft, M.P., and M.C.; Writing – Review & Editing, M.P., M.D.S., B.M., A.B.C., E.G.G, B.C. and M.C.; Funding Acquisition, E.G.G. and M.C.; Supervision, E.G.G. and M.C.

